# Alzheimer’s genetic risk effects on cerebral blood flow are spatially consistent and proximal to gene expression across the lifespan

**DOI:** 10.1101/2020.12.31.424949

**Authors:** H.L Chandler, R.G Wise, D.E Linden, J Williams, K Murphy, T.M Lancaster, Alzheimer’s Disease Neuroimaging Initiative (ADNI)

**Author notes:** Corresponding author: Dr Hannah L Chandler, Department: Cardiff University Brain Research Imaging Centre, School of Psychology, Cardiff, University, UK.

## Abstract

Cerebrovascular dysregulation is a hallmark feature of Alzheimer’s disease (AD), where alterations in cerebral blood flow (CBF) are observed decades prior to symptom onset. Genome-wide association studies (GWAS) show that AD has a polygenic aetiology, providing a tool for studying AD susceptibility across the lifespan. Here, we ascertain whether AD genetic risk effects on CBF previously observed (Chandler et al., 2019) remain consistent across the lifespan. We further provide a causal mechanism to AD genetic risk scores (AD-GRS) effects by establishing spatial convergence between AD-GRS associated regional reductions in CBF and mRNA expression of the proximal AD transcripts using independent data from the Allen Brain Atlas. We analysed grey matter (GM) CBF in a young cohort (N=75; aged 18-35) and an older cohort (N=90; aged 55-85). Critically, we observed that AD-GRS was negatively associated with whole brain GM CBF in the older cohort (standardised β −0.38 [−0.68 – −0.09], P = 0.012), consistent with our prior observation in younger healthy adults (Chandler et al., 2019). We then demonstrate that the regional impact of AD-GRS on GM CBF was spatially consistent across the younger and older samples (r = 0.233, P = 0.035). Finally, we show that CBF across the cortex was related to the regional expression of the genes proximal to SNP’s used to estimate AD-GRS in both younger and older cohorts (Z_TWO-TAILED_ = −1.99, P= 0.047; Z_TWO-TAILED_ = −2.153 P = 0.032, respectively). These observations collectively demonstrate that AD risk alleles have a negative influence on brain vascular function and likely contribute to cerebrovascular changes preceding the onset of clinical symptoms, potentially driven by regional expression of proximal AD risk genes across the brain. Our observations suggest that reduced CBF is an early antecedent of AD and a key modifiable target for therapeutic intervention in individuals with a higher cumulative genetic risk for AD. This study will further enable identification of key molecular processes that underpin AD genetic risk related reductions in CBF that could be targeted decades prior to the onset of neurodegeneration.

## Introduction

Variability of cerebrovascular function is heritable and partly explained by additive effects of genetic factors that converge across several neurobiological processes (Ikram et al., 2018). In Alzheimer’s disease (AD), cerebrovascular dysregulation is a key concomitant factor (Kelleher & Soiza, 2013), and is one of the earliest markers of AD pathophysiology (Iturria-Medina et al., 2016; Kelleher & Soiza, 2013). Decreases in cerebrovascular function are observed both in patients with AD and young individuals with an increased risk of dementia (Chandler et al., 2019; Filippini et al., 2011; Montagne et al., 2020; Wolters et al., 2017) This broadly suggests that altered cerebrovascular function is a risk factor for AD, rather than a consequence of the disease, which may be present across an individual’s lifespan.

Genome-wide association studies (GWAS) demonstrate that AD is also highly polygenic, where potentially thousands of common risk alleles confer susceptibility for disease (Kunkle et al., 2019). Although polygenic analysis has shown utility in predicting AD (Escott-Price, Shoai, Pither, Williams, & Hardy, 2017; Escott-Price et al., 2015), the neurobiological mechanisms by which these loci confer risk remains poorly understood, particularly in relation to cerebrovascular function. Furthermore, the impact of these risk alleles across the lifespan has been seldom explored. Several studies have suggested that the influence of AD risk alleles may be age-dependent (Matura et al., 2016), while other large studies demonstrate that the impact of AD risk alleles on risk factors such as cognition are influential across the entire lifespan (Hill et al., 2016). However, the impact of AD risk alleles on *in-vivo* measures of brain function has not been investigated across the lifespan.

In our previous work we used arterial spin labelling (ASL) with MRI to quantify regional cerebral perfusion in young healthy individuals (18-35 years) and observed negative associations between AD-polygenic risk and regional perfusion, as well as lower CBF in those who possess a copy of the *APOE*-ε4 allele. Our findings suggest that vascular alterations in those with a broad increased genetic risk for AD manifest decades prior to symptom onset (Chandler et al., 2019). While our prior work provided insight into the influence of genetic risk factors on the cerebrovasculature in early adulthood, it is not yet known whether the influence of AD genetic risk scores on GM CBF remains consistent across the lifespan.

In the current study, we aim to determine the impact of AD risk alleles on CBF in an older population (mean age= 70). We anticipate that the combined influence of AD risk alleles will be associated with a reduction in global CBF (similar to our findings in Chandler et al., 2019). Here, one predicts that either i) the effects of AD risk alleles on CBF remain consistent or ii) demonstrate a more pronounced influence later in life. As AD risk alleles are likely to confer susceptibility by influencing expression of proximal genes, we further anticipate that regional CBF is spatially related to the expression of these AD risk alleles. In order to address this hypothesis, we probe the Allen Human Brain Atlas (AHBA) to understand the relationship between AD risk gene expression and regional CBF across the cortex to determine if the influence of AD risk alleles can be explained by the regional co-expression of gene transcripts proximal to these AD risk loci (Arnatkevic lute, Fulcher, & Fornito, 2019). Specifically, we sought to investigate whether brain-wide AD-related gene expression covaries with regional variation in CBF. These analyses will establish the regional cortical co-expression of AD risk genes and AD-risk gene related CBF reductions, providing a plausible mechanistic link between AD risk loci and a well-established pathophysiological process in AD.

## Methods

### Participants

#### ADNI cohort

A total of ninety participants, classified as either healthy controls (N = 30) or having mild cognitive impairment (MCI, N = 60) took part in a series of MRI scans as part of their involvement in the ADNI protocol. Participants were removed if they also contributed to IGAP AD GWAS (N=2). Some participants were scanned at several timepoints, where the final number of discrete data points N_OBSERVATIONS_ = 131, where N_PARTICIPANTS_ = 48 completed one scan and N_PARTICIPANTS_ = 42 completed more than one scan for the final analysis. See Table 1 for further demographic information.

**Table 1.** MCI = Mild cognitive impairment; Education = years; ICV = Intracranial Volume.

| | Combined Sample<br>N=90 | | APOE ε4 (-)<br>N=62 | | APOE ε4 (+)<br>N=28 | | P |
| --- | --- | --- | --- | --- | --- | --- | --- |
| Sex | M=40 / F=50 |  | M=35 / F=27 |  | M= 15 / F=13 |  | .979 |
| MCI | M = 30 / F = 60 |  | M = 20 / F = 42 |  | M = 10 / F = 18 |  | .935 |
|  | M | SD | M | SD | M | SD |  |
| Age | 70.25 | 6.50 | 70.33 | 6.53 | 70.08 | 6.55 | .866 |
| Education | 16.68 | 2.59 | 16.94 | 2.63 | 16.11 | 2.44 | .151 |
| ICV | 1.5E+06 | 1.4E+05 | 1.5E+06 | 1.4E+05 | 1.5E+06 | 1.7E+05 | .682 |

#### Cardiff cohort

Our younger sample was identical to our previous sample (Chandler et al., 2019) and consisted of seventy-five (N_FEMALE_ = 47), right-handed individuals of western European descent, aged between 19-35. For further sample characterisation (including ethics, exclusion criteria, genotyping methods, see Chandler et al., (2019).

### Creation of polygenic scores

Polygenic score calculations were performed according to the procedure described by the International Schizophrenia Consortium, using the --score command in PLINK, via a wrapper function provided in the PRSice v1.25 software package (Euesden, Lewis, & O’Reilly, 2015). Training data were from a recent AD GWAS (Kunkle et al., 2019), where SNPs were removed from summary statistics / genotype data if they had a low minor allele frequency (P < 0.01) and data were pruned for linkage disequilibrium removing SNPs within 500 kb and *R2* > 0.1 with a more significantly associated SNP. For the creation of the AD-GRS, we considered SNPs that were associated with AD that surpassed the GWAS threshold (PT < 5 × 10-8), as performed and to make comparable to our original study (Chandler et al., 2019). Individual *APOE* ε4 status was independently modelled in all analyses. Twenty-six SNPs were considered in the final AD-GRS calculation (see Figure 2). To minimise potential confounding from population stratification linked to AD-GRS we included the first five principle components from a linkage-disequilibrium (LD) pruned version of the genotypes as covariates in all analysis (Choi, Mak, & O’Reilly, 2020).

### Imaging procedures and analysis of CBF

#### ADNI cohort

A 3T siemens PICORE MRI sequence (Wong, Buxton, & Frank, 1997) with pulsed ASL (or Q2TIPS) (Luh, Wong, Bandettini, & Hyde, 1999). The sequence parameters include repetition time (TR) = 3400 ms, echo time (TE) = 12 ms, TI1 = 700 ms, TI2 = 1900 ms, field of view (FOV) = 256 mm × 256 mm, number of slices: 24 axial, slice thickness = 4 mm, and image matrix size = 64 × 64. Pre-processing steps were conducted in SPM8 and included motion correction of individual ASL frames by rigid body transformation and least squares fitting. To obtain perfusion weighted images, the ASL data were then split into tag and control images and the mean-untagged data were subtracted from the mean-tagged data. The first volume of the ASL scan was used in place of an M0 (providing fully relaxed signal) to estimate blood-water-density proxy and used for calibration. A 3D MPRAGE T1-weighted sequence was collected for registration with the following parameters: TR = 2300ms, TE = 2.98ms, TI =900ms, 176 sagittal slices, FOV = 256×240mm^2^, voxel size=1.1×1.1×1.2mm^3^, flip angle=9°. The perfusion data were registered to T1 space and rescaled to obtain CBF in ml/100g/min. For both cohorts, GM CBF values were sampled in native space across 82 cortical and subcortical parcellations as segmented using a FreeSurfer template (Desikan et al., 2006; Potvin, Dieumegarde, Duchesne, & Alzheimer’s Disease Neuroimaging, 2017). Full analysis including details of distortion correction, registration and partial volume correction can be found via the ADNI web page (http://adni.loni.usc.edu).

#### Cardiff cohort

Imaging data were collected on a 3T General Electric (GE) MRI scanner. Anatomical T1-weighted images were acquired with a 3D fast spoiled gradient echo sequence (FSPGR). Sequence parameters included: 172 contiguous sagittal slices with a slice thickness of 1 mm, TR = 7.9, TE = 3ms, inversion time of 450ms, flip angle = 20°, a FOV of 256 × 256 × 176 mm, matrix size 256 × 256 × 192 to yield 1 mm isotropic voxel resolution images. Resting CBF data were collected using a pseudo-continuous arterial spin labelling (PCASL) sequence. The study consisted of a single MRI session (which also comprised other functional and structural scans), and the PCASL sequence that lasted approximately 6 minutes. A PCASL sequence was acquired and included a 3D fast spin echo (FSE) spiral multi-slice readout. The sequence parameters included: number of excitations = 3, time to echo=32ms, echo time train length=64, TR=5.5seconds, matrix size=48×64×60, FOV=18×23×18cm, tag= 1500ms, PLD = 1500ms. For detail regarding ASL pre-processing pipeline see (Chandler et al., 2019).

### Gene expression analysis

Publicly available human gene expression data from six post-mortem donors (N_FEMALE_ = 1), aged 24-57 (42.5±13.38) were obtained from the Allen Institute (Hawrylycz et al., 2012). Data reflect the microarray normalization pipeline implemented in March 2013 (http://human.brain-map.org) and analyses were conducted according to the guidelines of the Yale University Human Subjects Committee. Normalised brain-wide gene transcript expression was mapped to eighty-two cortical /subcortical regions of interest as defined by the Desikan-Killiany atlas in abagen v0.0.3 (Arnatkevic lute et al., 2019) available to download at https://zenodo.org/record/3688800-.XnoejVKcYWo.

### Statistical analysis

To maximise consistency of regression models across both samples, we included the same covariates for both cohort analyses. Predictors were regressed against i) whole GM CBF and ii) regional GM CBF for the eighty-two cortical / subcortical regions as defined by the Desikan-Killiany-Tourville (DKT) Atlas (Potvin et al., 2017). The fixed effects of AD-GRS (P_T_ < 5 × 10^−8^) and *APOE* (modelled via the absence / presence (0 / 1) of an ε4 allele) were modelled while controlling for age, biological sex, education, ICV and the first five genetic principal components, acquired via the LD-pruned datasets. For the ADNI cohort, we further included fixed effect covariates for i) diagnostic status (healthy control / mild cognitive impairment) ii) years of education and iii) site and random effects for iv) visit code and v) subject, modelled as repeated measures. We employed outlier labelling / detection (Hoaglin & Iglewicz, 1987) to dynamically remove data points for each GM CBF dependent variable to minimize the impact of outlier data points. In order to control for false positives for each regional gene expression analysis, we compare each correlation to i) 10,000 randomly generated regional gene expression profiles equalling N_UNIQUE-GENES_ = 46 and ii) randomly generated brain-wide regional ‘phenotypes’ where we simulate 10,000 normal distributions for eighty-two effect sizes scaled to either CBF or AD-GRS effects on CBF. In order to assume that the strength of the gene expression – CBF covariation is more pronounced than expected by chance, the observed z-transformed correlation must surpass the alpha tail (Z > 1.96 / 95% CI: two-tailed) for both (simulated gene expression and brain-wide CBF / AD-GRS effects) simulated distributions.

## Results

### 3.1 Cerebral blood flow across the lifespan

First, we observed that regional GM CBF showed a consistent pattern of positive covariation between the younger and older cohorts (Figure 1A-B), where cortical regions that showed higher perfusion (ml/100g/min) in the younger sample was also comparably higher in the older sample (r = 0.328: P = 0.002; Figure 1C), suggesting a pattern of consistent, regional variation in GM CBF across the lifespan.

**Figure 1.**
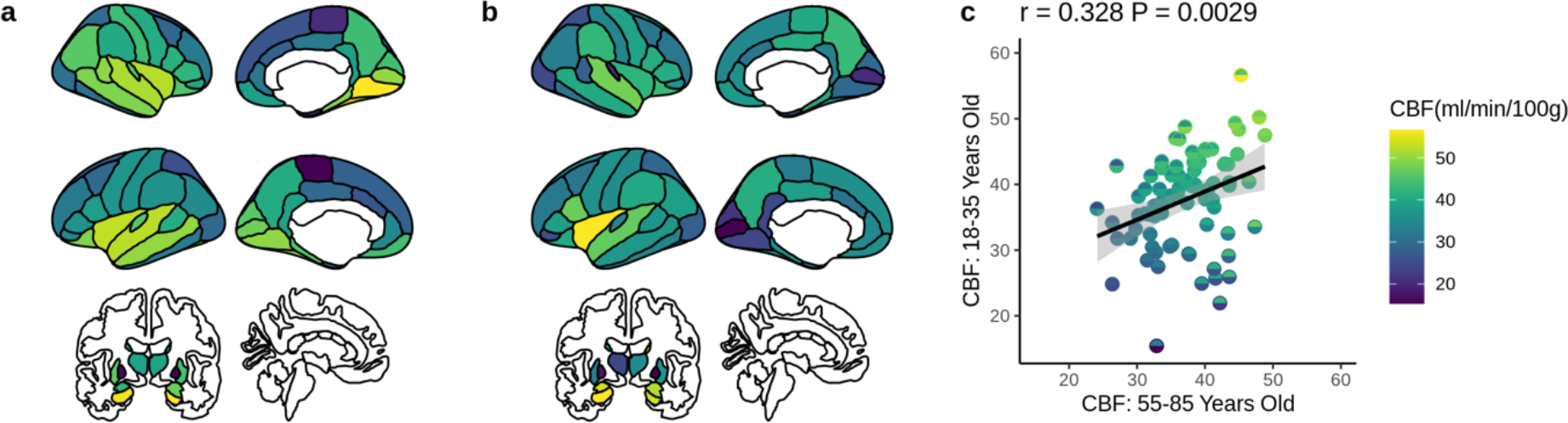
Regional GM CBF (ml/100g/min) in a) the younger cohort (aged: 18-35) previously described in Chandler et al., 2019 and b) an older cohort (aged: 55-85) and c) Regional GM CBF (ml/100g/min) comparison for young (y-axis) and old (x-axis) across all eighty-two cortical / subcortical regions.

### 3.2. AD-GRS effects on whole brain cerebral blood flow (ml/min/100g)

Similar to our original discovery (Chandler et al., 2019), we observed a significant negative association between whole brain GM CBF and AD-GRS in the older (55-85 years) ADNI sample (β = −0.38; P = 0.012) after controlling for all covariates. Unlike our observation in younger individuals (Chandler et al., 2019) we did not observe a significant association between *APOE* ε4 absence / presence and whole brain GM CBF in the older cohort (β = 0.41; P = 0.177). For all fixed effects and confidence intervals observed in the whole brain GM CBF analysis, see Table 2.

**Table 2.**
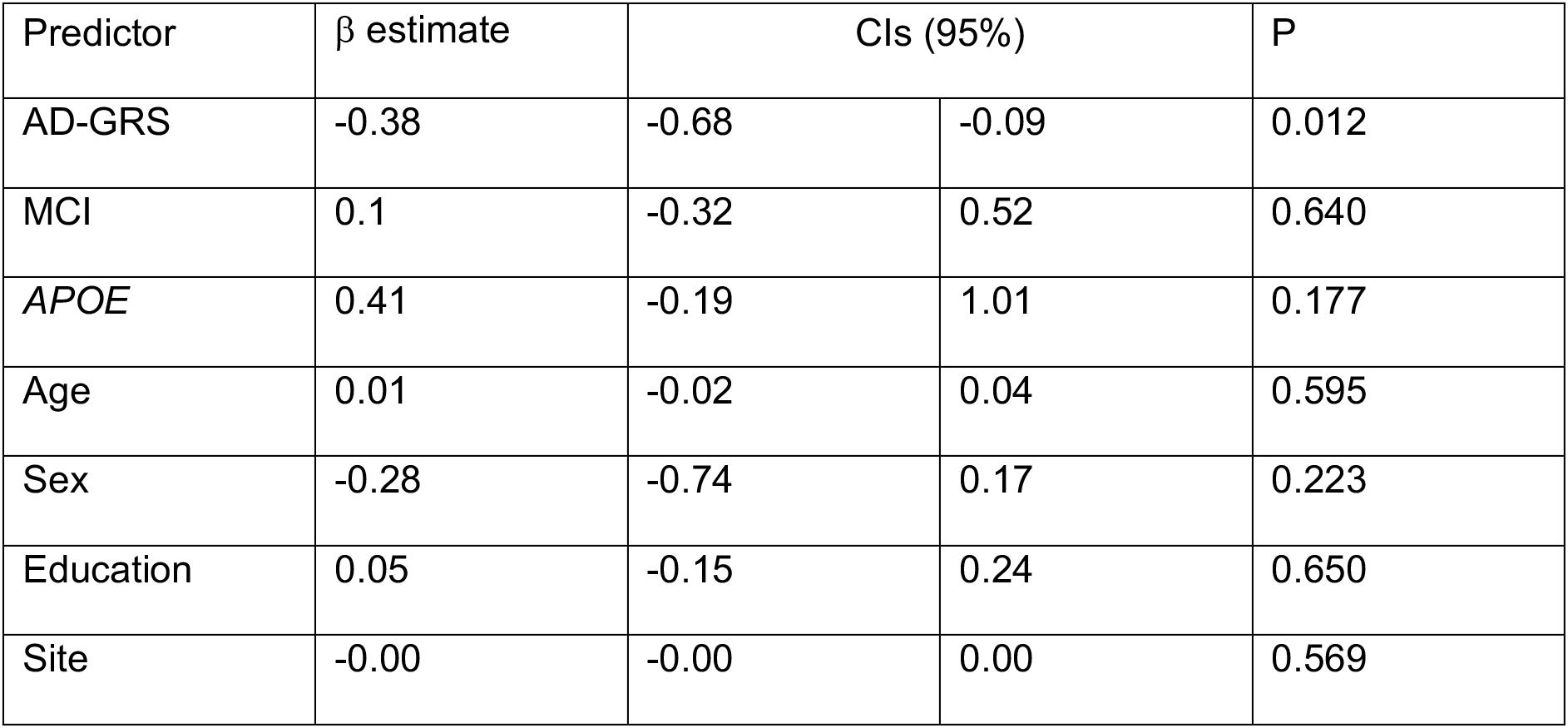
Fixed effect predictors (β estimate and 95% confidence intervals) regressed against whole brain GM CBF in the final sample of ADNI participants controlling for the top five principal components (PCs) as additional covariates of no interest and visit code / subject as random effects. MCI = mild cognitive impairment.

In order to assess the impact of each of the twenty-six SNPs in our AD-GRS model, we performed a linear regression analysis where each individual SNP was regressed in an additive model against whole brain GM CBF, controlling for all aforementioned covariates. Consistent with broad polygenic modelling assumptions, we observed a general propensity for SNPs that increase risk for AD (odds ratio (OR) > 1) to associate with reduced whole GM CBF, while alleles that conferred relative protection (OR < 1) for AD where associated with an increase in whole GM CBF (Figure 2; sign test for direction of effects: P = 0.002; independent samples t-test for mean risk / protective beta estimates: t = 2.55, P = 0.018).

**Figure 2.**
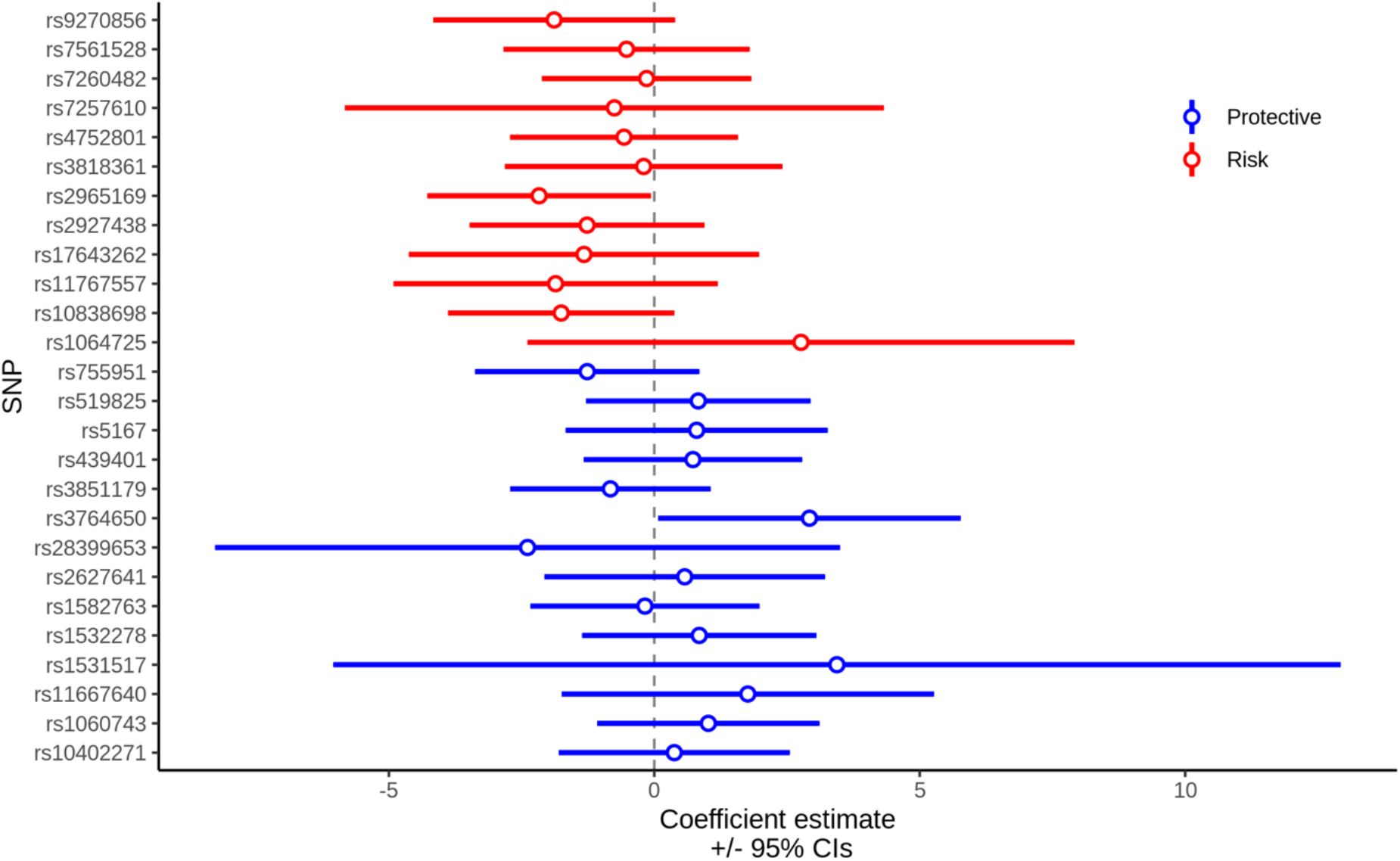
Diagnostic plot, demonstrating individual effects of AD risk (red) and protective (blue) SNPs on whole brain GM CBF, controlling for covariates in the older sample (55-85 years old). Circles / lines represent adjusted effect sizes and 95% confidence intervals.

### 3.3. Comparing AD-GRS effects on cerebral blood flow in early adulthood and older age

As we observed an association between AD-GRS and whole brain GM CBF for both younger and older samples (Figure 3a), we proceeded to explore the association at a regional level. We repeated the linear mixed-model analysis across eighty-two cortical / subcortical regions. Building upon our initial analysis in the younger cohort (Chandler et al., 2019, replotted here in Figure 3b), we observed a significant relationship between regional effect sizes across the brain, where the most / least pronounced effects of AD-GRS were comparable between young and older samples (Figure 3d). However, in the ADNI sample of older individuals (55-85 years old) we found no specific regions with significant effects after correcting for false discovery rate (lowest P_UNCORRECTED_ = 0.006; left hemisphere, insula), We did not observe the influence of *APOE* ε4 status on a) whole brain GM CBF in the older sample and b) a regional effect of *APOE* ε4 on the younger sample, so did not proceed to investigate similarity between samples for *APOE* ε4 GM CBF effects at a regional level.

**Figure 3.**
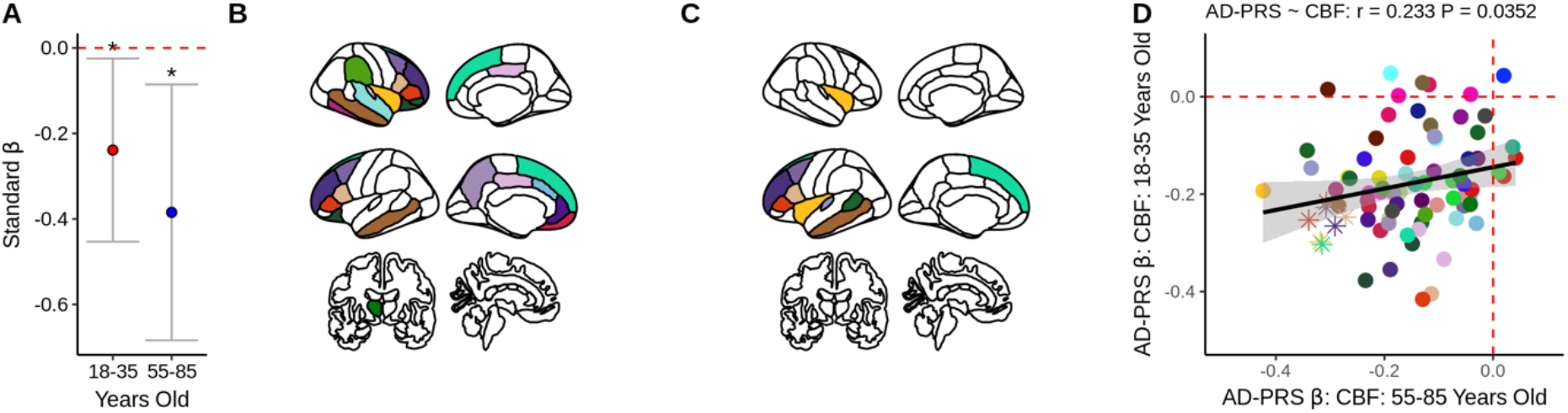
Standardised AD-GRS effects on A) whole brain GM CBF for the young (18-35) and older (55-85) cohorts; * indicates P < 0.05, error bars represent 95% confidence intervals. Regional influence of AD-GRS on b) the young cohort (previously described in Chandler et al., 2019) and c) the ADNI cohort, both P_UNCORRECTED_ < 0.05, for visual purposes. D) Linear relationship of effect sizes across the brain when comparing standardised beta estimates for all cortical regions between sample B & C, where data points represented as asterisk reflect P < 0.05 in both samples. Each point in the scatter plot represents one cortical / subcortical region.

### 3.4. Regional AD risk gene expression overlap

We calculated the average transcript expression of AD risk genes proximal to the twenty-six SNPs used in AD-GRS model (N_UNIQUE-GENES_ =46) for the eighty-two cortical / subcortical regions (Figure 4A). We then correlated regional mean AD risk gene expression with 1) regional CBF (ml/100g/min) for the younger and older samples. We observed that mean AD risk gene expression was negatively associated with regional GM CBF in the young (Z = - 1.99, P = 0.047) and the older sample (Z = −2.153, P = 0.031), suggesting that AD risk gene expression is highest in cortical regions where GM CBF is generally lower, regardless of AD genetic risk.

**Figure 4.**
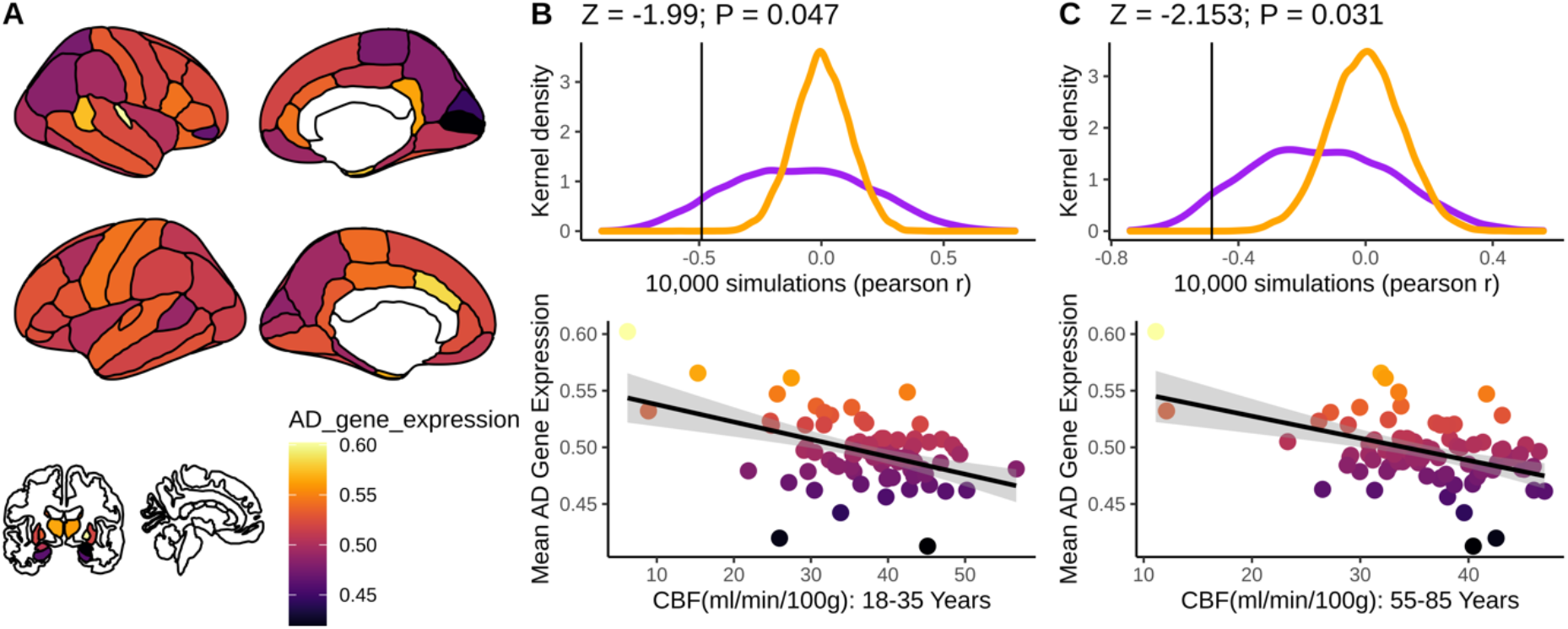
A) Mean, normalised AD risk gene expression across eighty-two cortical / subcortical regions. B-C (lower) Scatter plots show relationship between regional mean AD gene expression (A) and regional CBF for the 18-35 year old (B) and 55-85 year old (C) cohorts. B-C (upper) Distribution of 10,000 simulated i) randomly selected mean gene expression profiles (purple curve) and ii) randomly simulated regional values (scaled to CBF range; orange curve). Solid black vertical lines represent the actual, observed correlation between mean AD gene expression (B-C, lower).

## Discussion

We sought to further investigate the impact of common AD genetic risk alleles on cerebral perfusion. Critically, the negative association we observed between AD-GRS and whole brain GM CBF in our prior work (Chandler et al., 2019) was also evident in the older population. This observation was further supported by evidence that regional effects of AD-GRS on GM CBF were correlated across samples. This suggests that the regional impact of AD-GRS on regional CBF remains consistent across the lifespan, and preferentially influences specific cortical structures previously implicated in preclinical models of AD related pathophysiology. Together these observations show that the cumulative impact of AD risk loci on this hallmark feature of AD pathogenesis is consistent across the lifespan.

The mechanisms by which common (e.g. intronic and intergenic) SNPs identified via GWAS confer susceptibility are largely unknown. However, a growing body of work suggests that these SNP’s act as expression quantitative trait locus (eQTLs) and influence the expression of AD risk genes. Here, we tested the hypothesis that the brain-wide CBF variability would be spatially convergent with the expression AD risk gene transcripts. We found consistent regional covariation between mean AD risk gene expression and regional CBF perfusion pattern in both young and old cohorts. These findings demonstrate that the impact of AD-GRS on perfusion may confer susceptibility via the altered expression of proximal AD genes.

Cerebral blood flow shows a gradual and steady decrease across the lifespan (Bertsch et al., 2009; Devous, Stokely, Chehabi, & Bonte, 1986; Hagstadius & Risberg, 1989; Heo et al., 2010; Lu et al., 2011). Here we showed spatial convergence of CBF variation between the young and old cohorts, suggesting that regional variability in GM CBF across the cortex remains largely consistent across age. It is not entirely understood why there is variability in GM CBF at rest across the brain. However, prior evidence has shown that brain perfusion closely correlates with brain function and metabolism (Detre, Wang, Wang, & Rao, 2009), suggesting that variability in regional perfusion may reflect differences in energy demand across the cortex at rest.

In the second analysis, we showed that the association between regional CBF and AD-GRS in the young cohort correlated positively with the association between regional CBF and AD-GRS in the older cohort. The most significant effects were mostly observed in the frontal and temporal cortical structures. Critically, this result demonstrates that the AD-GRS effects seen in the older cohort are regionally congruent with those in the younger cohort. Our findings suggest that SNPs included in the AD genetic/polygenic risk model have consistent negative effects on cerebral perfusion from young adulthood and throughout the lifespan. In addition to AD-GRS we also investigated the effects of *APOE* on CBF across the cohorts. We saw no influence of *APOE* ε4 status on whole brain or regional CBF in the ADNI cohort (unlike in our prior study (Chandler et al., 2019). We suggest that while the AD-GRS influence on CBF across the lifespan remains consistent, *APOE* status may have a more dynamic role in shaping CBF (Wierenga et al., 2013) and requires further investigation.

In our third analysis we used gene expression data to identify how AD risk genes expression across the cortex correlates with regional CBF. Our results show a negative association between AD gene expression and regional CBF, suggesting that AD risk genes may spatially covary with regional cerebral perfusion. Moreover, our findings demonstrate that the regional covariation between cerebral perfusion and AD gene expression occurs throughout the lifespan.

While amyloid and tau-genic hypotheses provide important insight into preclinical AD models (Bloom, 2014; Gotz, Chen, van Dorpe, & Nitsch, 2001; Ittner & Gotz, 2011; Lewis et al., 2001), vascular dysregulation occurs prior to this AD pathophysiology (Iturria-Medina et al., 2016). We provide additional support for vascular dysregulation and hypoperfusion as early markers of AD risk that may be observed during young adulthood. Moreover, we suggest that cerebral perfusion is a potentially important AD related pathological feature and should be considered as a target for therapeutic intervention.

Our observations should be considered with the following limitations. First, while we observed consistent AD-GRS effects across the lifespan, we did not observe *APOE* related effects in our older sample. This may be explained by dynamic *APOE* effects that have recently been discovered in recent structural MRI studies (Brouwer et al., 2020) or lower statistical power afforded in smaller samples. Third, while we aimed to generate a single regional metric of AD gene expression, AD risk SNPs may confer risk by increasing or decreasing proximal gene expression, so generating regional expression profiles that consider this variation (rather than using average AD gene expression) will be useful in refining the impact of AD risk SNPs on regional gene expression,

To conclude, we demonstrate a consistent negative influence of additive genetic AD risk on cerebral perfusion across the lifespan, which was also related to regional expression of proximal AD risk genes across the cortex. Thus, reduced CBF may be a central, and proximal, process in the pathophysiology of AD, and a mechanism by which AD risk genes exert their adverse effects on brain structure and function.

## ACKNOWLEDGEMENTS

HLC is funded by a Wellcome Strategic Award [104943/Z/14/Z].

KM is funded by the Wellcome Trust [WT200804].

TML acknowledges funding via a Sêr Cyrmu II Fellowship (East Wales European Regional Development Funds (PNU-80762-CU-14).

We are grateful to all professionals, patients and volunteers involved with the National Centre for Mental Health (NCMH) who took part in our Cardiff study. NCMH is funded by the National Institute for Social Care and Health Research (NISCHR), Welsh Government, Wales (Grant No. BR09).

Data collection and sharing for this project was funded by the Alzheimer’s Disease Neuroimaging Initiative (ADNI) (National Institutes of Health Grant U01 AG024904) and DOD ADNI (Department of Defense award number W81XWH-12-2-0012). ADNI is funded by the National Institute on Aging, the National Institute of Biomedical Imaging and Bioengineering, and through generous contributions from the following: AbbVie, Alzheimer’s Association; Alzheimer’s Drug Discovery Foundation; Araclon Biotech; BioClinica, Inc.; Biogen; Bristol-Myers Squibb Company; CereSpir, Inc.; Cogstate; Eisai Inc.; Elan Pharmaceuticals, Inc.; Eli Lilly and Company; EuroImmun; F. Hoffmann-La Roche Ltd and its affiliated company Genentech, Inc.; Fujirebio; GE Healthcare; IXICO Ltd.; Janssen Alzheimer Immunotherapy Research & Development, LLC.; Johnson & Johnson Pharmaceutical Research & Development LLC.; Lumosity; Lundbeck; Merck & Co., Inc.; Meso Scale Diagnostics, LLC.; NeuroRx Research; Neurotrack Technologies; Novartis Pharmaceuticals Corporation; Pfizer Inc.; Piramal Imaging; Servier; Takeda Pharmaceutical Company; and Transition Therapeutics. The Canadian Institutes of Health Research is providing funds to support ADNI clinical sites in Canada. Private sector contributions are facilitated by the Foundation for the National Institutes of Health (http://www.fnih.org). The grantee organization is the Northern California Institute for Research and Education, and the study is coordinated by the Alzheimer’s Therapeutic Research Institute at the University of Southern California. ADNI data are disseminated by the Laboratory for Neuro Imaging at the University of Southern California.

